# Accurate PROTAC targeted degradation prediction with DegradeMaster

**DOI:** 10.1101/2025.02.03.636343

**Authors:** Jie Liu, Michael Roy, Luke Isbel, Fuyi Li

**Affiliations:** South Australian Immunogenomics Cancer Institute (SAiGENCI), The University of Adelaide, Adelaide, 5005, South Australia, Australia; Adelaide Centre for Epigenetics, The University of Adelaide,Adelaide, 5005, South Australia, Australia

## Abstract

**Motivation:** Proteolysis-targeting chimeras (PROTACs) are heterobifunctional molecules that can degrade ‘undruggable’ protein of interest (POI) by recruiting E3 ligases and hijacking the ubiquitin-proteasome system. Some efforts have been made to develop deep learning-based approaches to predict the degradation ability of a given PROTAC. However, existing deep learning methods either simplify proteins and PROTACs as 2D graphs by disregarding crucial 3D spatial information or exclusively rely on limited labels for supervised learning without considering the abundant information from unlabeled data. Nevertheless, considering the potential to accelerate drug discovery, it is critical to develop more accurate computational methods for PROTAC-targeted protein degradation prediction.

**Results:** This study proposes DegradeMaster, a semi-supervised E(3)-equivariant graph neural network-based predictor for targeted degradation prediction of PROTACs. DegradeMaster leverages an E(3)-equivariant graph encoder to incorporate 3D geometric constraints into the molecular representations and utilises a memory-based pseudo-labeling strategy to enrich annotated data during training. A mutual attention pooling module is also designed for interpretable graph representation. Experiments on both supervised and semi-supervised PROTAC datasets demonstrate that DegradeMaster outperforms state-of-the-art baselines, with substantial improvement of AUROC by 10.5%. Case studies show DegradeMaster achieves 88.33% and 77.78% accuracy in predicting the degradability of VZ185 candidates on BRD9 and ACBI3 on KRAS mutants. Visualization of attention weights on 3D molecule graph demonstrates that DegradeMaster recognises linking and binding regions of warhead and E3 ligands and emphasizes the importance of structural information in these areas for degradation prediction. Together, this shows the potential for cutting-edge tools to highlight functional PROTAC components, thereby accelerating novel compound generation.

**Availability:** The source code and datasets are available at https://github.com/Jackson117/DegradeMaster and https://zenodo.org/records/14715718.

## 1. Introduction

Proteolysis-targeting chimeras (PROTACs) are heterobifunctional molecules comprising a target protein-of-interest (POI) binding moiety (warhead), a linker, and an E3 ubiquitin ligase binding moiety [1, 2]. By forming a ternary POI-PROTAC-E3 ligase complex [3, 4], PROTACs exploit the ubiquitin-proteasome system to induce polyubiquitination and subsequent proteasomal degradation of the target protein [5]. As an emerging therapeutic strategy, PROTACs offer advantages over traditional small-molecule inhibitors that are attractive to target previously “undruggable” proteins [6], including addressing non-catalytic functions of target proteins, potential for enhanced selectivity and specificity, and preventing the accumulation of drug targets [7, 8].

Currently, PROTAC development is highly dependent on iterative strategies for design, synthesis, biological evaluation, and optimization [9, 10], which can be time-intensive and costly. Thanks to the increasing availability of detailed and high-quality PROTAC data [11, 12, 13] and extensive PROTAC databases like PROTAC-DB [14], machine learning has emerged as a viable tool for predicting PROTAC degradation. Traditional machine learning models, such as Support Vector Machine (SVM) [15] and Random Forest (RF) [16], leverage physicochemical descriptors of PROTAC molecules and target proteins for predictive modelling [17, 18]. Recent research has advanced the use of graphs to model the complex structural intricacies of the PROTAC system. Deep learning approaches, including Graph Neural Networks (GNNs) [19, 20], Long Short-Term Memory networks (LSTMs) [21] and Transformers [22], have been employed to encode both structural and attribute-based information of PROTAC molecules and target proteins, facilitating downstream tasks such as degradation prediction [23, 24, 25].

Despite their notable performance, deep learning models face two major limitations: First, existing deep learning approaches [25, 24] represent proteins and PROTACs as 2D graphs for simplicity, disregarding their 3D atomic coordinates.

However, the 3D spatial arrangement of atoms is crucial for determining binding interactions and molecular geometry, which directly influence the efficacy of PROTAC-mediated target protein degradation. The omission of 3D spatial information leads to the loss of structural compatibility, thereby diminishing model accuracy. Second, existing predictive methods exclusively rely on supervised learning, yet the datasets from PROTAC-DB are predominantly unlabeled, with over 80% of the data lacking annotation of degradation activity (i.e., lacking values for *DC*_50_ and *Dmax*; respectively the half-maximal degradation concentration and percentage maximum degradation achieved). This scarcity of labelled data increases the risk of overfitting, particularly when employing complex model architectures. Consequently, developing a semi-supervised learning approach that incorporates the abundant unlabeled data into the training process offers a promising solution to mitigate the issue of limited annotations and improve prediction accuracy.

To address these challenges, we propose a semi-supervised E(3)-equivariant graph neural network (DegradeMaster) for PROTAC degradation prediction. DegradeMaster integrates five key components: (1) a 3D molecule graph construction module to capture spatial and structural information from the PROTAC system, (2) an E(3)-equivariant encoder to incorporate geometric constraints such as distances and angles into the molecular representations, (3) a feature selection module to extract informative chemical descriptors as attribute features, (4) a mutual attention pooling module to compute comprehensive graph representations, and (5) a memory-based pseudo-labeling module to expand the size of annotated data during training. Benchmarking experiments on two PROTAC-DB dataset variants demonstrate that DegradeMaster outperforms PROTAC-STAN [24] and other state-of-the-art models [25] for PROTAC degradation prediction.

## 2. Materials and Methods

### 2.1. Data collection and preprocessing

To evaluate the performance of our model and baseline methods, we utilised data from the PROTAC-DB 3.0 database [14]. The latest release of PROTAC-DB 3.0 comprises 9,380 PROTAC entries, including 569 warheads, 107 E3 ligands, and 5,753 linkers. Each entry includes detailed information such as the PROTAC’s SMILES representation, and UniProt ID of the POI and the E3 ligase. Additionally, the database provides data on PROTAC degradation activity, including *DC*_50_ and *Dmax*. These metrics serve as standard measures of a PROTAC’s efficiency in degrading target proteins [7, 4], where a lower *DC*_50_ and a higher *Dmax* indicate relatively high degradation potential.

We first removed entries that lack critical information, e.g., the UniProt ID of the POI or E3 ligase. For degradation labels, following [25], we utilized both explicit *DC*_50_/*D*_*max*_ values and implicit values inferred from experimental descriptions to predict PROTAC degradation activity. A PROTAC is considered to have low degradation activity if *DC*_50_ is greater than or equal to 100 nM and *Dmax* is below 80%, otherwise it is labelled with high degradation activity. Crystal structures of POIs and E3 ligases are sourced from the Protein Data Bank (PDB) [26], while proteins without available crystal structures are supplemented with predicted structures generated by AlphaFold 2 [27]. We applied Smina [28] to dock the warhead and E3 ligand to POI and E3 ligase, respectively, defining the residues within 5 Å of the ligand as the pocket. Using these criteria, we constructed a supervised PROTAC dataset consisting of 620 high-activity entries and 1,011 low-activity entries. Additionally, we curated a semi-supervised PROTAC dataset containing 8,603 entries in total, incorporating the same labelled subset as the supervised dataset. Dataset information is elaborated in Table 1.

**Table 1.** Statistics of the two PROTAC datasets extracted from PROTAC-DB 3.0 [14]. High and low indicate high and low degradation activities of PROTACs, respectively.

| Datasets | Entries | Warheads | E3-ligands | Linkers | High | Low |
| --- | --- | --- | --- | --- | --- | --- |
| PROTAC-1K | 1,503 | 360 | 80 | 1,500 | 577 | 925 |
| PROTAC-8K | 8,636 | 569 | 107 | 5,753 | 577 | 925 |

### 2.2. The framework of DegradeMasters

An overview of the architecture of DegradeMaster, and its key components are illustrated in Figure 1. As depicted in Figure 1 (A), DegradeMaster employs a multi-step framework designed to predict the degradation efficacy of PROTAC molecules by effectively capturing spatial, structural, and physicochemical information. It consists of five major components: 3D molecule graph construction, E(3) equivariant encoder, feature selection, mutual attention pooling, and label enrichment. The method begins by constructing 3D molecule graphs for the PROTAC, POI, and E3 ligase. Each graph represents atoms as nodes and chemical bonds as edges while incorporating intra- and inter-residue connections for proteins. 3D coordinates of each atom are recorded.

**Fig. 1.**
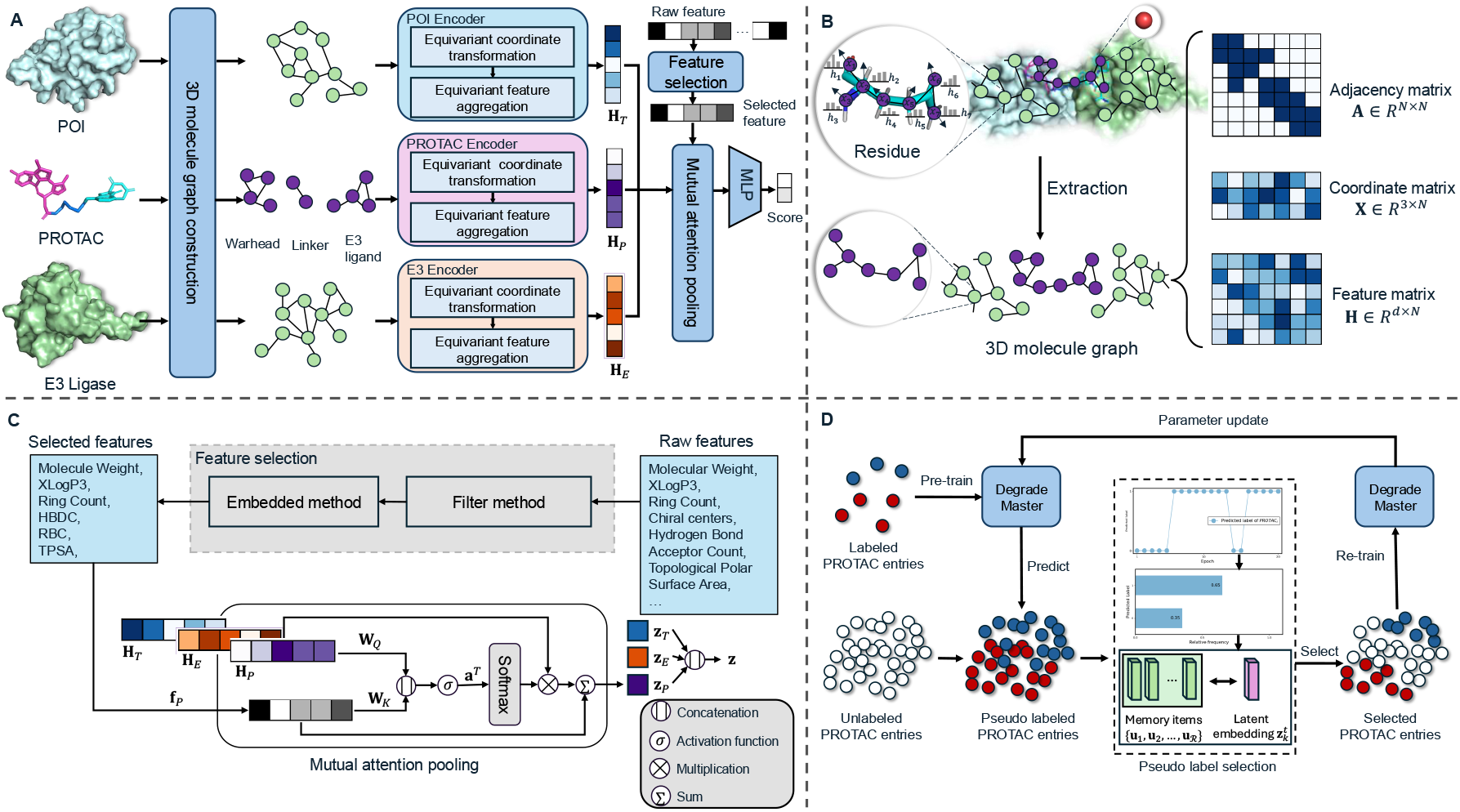
(**A**) Overall framework of DegradeMaster. (**B**) 3D molecule graph construction procedure. (**C**) Feature selection and mutual attention pooling. Abbreviation: Hydrogen Bond Donor Count (HBDC), Rotatable Bond Count (RBC), Topological Polar Surface Area (TPSA). (**D**) Memory enhanced pseudo-labeling.

Next, the E(3) equivariant encoder ensures the model respects transformations (rotation, reflection, and translation) in 3D space, which is crucial for maintaining consistency in spatial relationships during graph encoding. The encoder first iteratively updates node coordinates using messages derived from neighboring nodes and edges, and then integrates information across edges to update node embeddings, ensuring E(3) equivariance is preserved throughout.

To refine the input data, a feature selection module identifies the most informative molecular properties, filtering out redundant attributes or those that do not influence the model greatly. A two-step process, combining statistical metrics such as variance and chi-square scores with an embedded method using Gradient Boosting Decision Trees [29], is applied. This results in a concise set of chemical descriptors that enhance the model’s predictive accuracy. A detailed explanation is provided in Section 1 of the supplementary material. Supplementary Tables S1 and S2 present the statistical metrics of all chemical properties.

The encoded graph representations are then aggregated into molecule-level embeddings using a novel mutual attention-pooling strategy. Finally, a label enrichment strategy addresses the scarcity of labelled data by leveraging semi-supervised learning. Pseudo-labels for unlabeled samples are generated using training dynamics, combining a disagreement-based score with a novel memory-based metric to assess pseudo-label reliability. The model iteratively refines its predictions, incorporating the enriched pseudo-label set into training to improve performance.

A detailed introduction to each key component of the methodology is elaborated in the following section.

#### 2.2.1. 3D molecule graph construction

In this section, we represent the spatial and structural information of molecules using graphs. As illustrated in the top-left of Figure 1 (B), a PROTAC molecule is modelled with its atoms as nodes and chemical bonds as edges, forming the node set 𝒱_*P*_ and the edge set ℰ_*P*_, respectively. Proteins, specifically the POI and E3 ligase, are divided into residues (depicted as green dots in Figure 1 (B)), with each residue comprising multiple atoms (shown as purple dots). The node sets for the POI and E3 ligase graphs are denoted as 𝒱_*T*_ and 𝒱_*E*_, respectively, where each node *υ* ∈ 𝒱_*T*_ or 𝒱_*E*_ corresponds to an atom. The edge sets are defined as ℰ_*ζ*_ = {ℰ_*ζ*,intra_, ℰ_*ζ*,inter_} for *ζ* ∈ {*T, E*}, where ℰ_·,intra_ and ℰ_·,inter_ represent intra-residue and inter-residue connections, respectively. For each molecule graph 𝒢_*ζ*_ comprising *N*_*ζ*_ atoms, the adjacency matrix is represented as 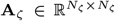, the feature matrix as 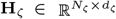 (where the *p*-th row of **H**_*ζ*_ encodes the feature vector of atom *p*), and the 3D atom coordinates as 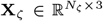. In summary, the constructed 3D molecule graph 𝒢_*ζ*_ is defined as 𝒢_*ζ*_ = {**A**_*ζ*_, **X**_*ζ*_, **H**_*ζ*_} for *ζ* ∈ {*P, T, E*}.

#### 2.2.2. (3) equivariance

In this section, we introduce the concept of E(3) equivariance. E(3) equivariance is a crucial inductive bias for predicting PROTAC degradation efficacy. Let **X** represent the coordinate matrix of the input 3D molecule graph 𝒢, and *T*_𝒢_ : **X** denote a linear transformation applied to **X**. A graph encoder *f*_*ϕ*_ : **X** → **Y** is defined as E(3) equivariant to 𝒢 if there exists a corresponding transformation *S*_𝒢_ : **Y** → **Y** in the output space such that the following condition holds:

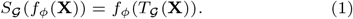

Specifically, let the linear transformation be *T*_𝒢_ (**X**) = **MX** + **B**, the equivalent transformation is *S*_𝒢_ (**Y**) = **MY** + **B**, where **M** ∈ ℝ^*N*×*N*^ denotes the rotation and reflection transition, while **B** ∈ ℝ^*N*×*N*^ denotes the translation transition. This ensures that the encoder output transforms consistently with the input under any E(3) transformation, preserving spatial and structural relationships critical for accurate degradation efficacy prediction.

#### 2.2.3. E(3) equivariant encoder

In this section, we elaborate on the design of our E(3) equivariant graph encoder. As shown in Figure 1 (A), the equivariant graph encoder consists of two key steps: equivariant coordinate transformation and equivariant feature aggregation. We adopt the equivariant graph neural network (EGNN) [30] as the base model in our coordinate update step. In the following part, we will introduce the two steps separately.

##### Equivariant coordinate transformation

Following the notion defined in section 2.2.1, given a 3D molecule graph 𝒢_*ζ*_ = {**A**_*ζ*_, **X**_*ζ*_, **H**_*ζ*_} for *ζ* ∈ {*P, T, E*}, each node *υ*_*i*_ ∈ *V*_*ζ*_ is associated with a coordinate vector **x**_*i*_ ∈ ℝ^3^ and a feature vector 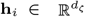. Let 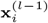 and 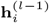 denote the coordinate embedding and feature embedding of node *υ*_*i*_ after (*l* − 1)-layer encoding, *e*_*ij*_ ∈ ℰ_*ζ*_ is the edges connecting to *υ*_*i*_, the coordinate vector 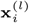 is calculated as follows:

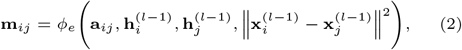

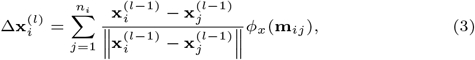

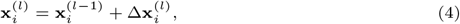

where **a**_*ij*_ is the edge attribute, *ϕ*_*e*_(·) and *ϕ*_*x*_(·) are MLPs that compute messages for edge and coordinate representations, respectively. *n*_*i*_ denotes the number of neighbours of node *υ*_*i*_.

##### Equivariant feature aggregation

The vector **m**_*ij*_ computed in Equation (3) indicates the message passed through edge *e*_*ij*_. Thus, the feature aggregation on node embedding 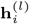 can be formed as:

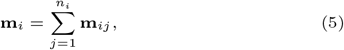

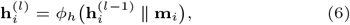

where *ϕ*_*h*_(·) is the Multilayer Perceptron (MLP) for node feature aggregation. By varying the input graphs among *𝒢*_*P*_, *𝒢*_*T*_, and *𝒢*_*E*_, we obtain node embeddings **H**_*P*_, **H**_*T*_, and **H**_*E*_ for PROTAC graph, POI graph, and E3 ligase graph in E(3)-equivariant manner, respectively.

#### 2.2.4. Mutal attention pooling

In Section 2.2.3, we incorporate the spatial and structural information of molecules and encode node embeddings in an E(3)-equivariant manner. Using the encoded node embeddings for the PROTAC molecule, 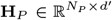, we apply a pooling operation to obtain the representation of the entire PROTAC graph. Unlike the mean pooling strategy employed in prior works [25], we introduce a mutual attention-based pooling strategy. This approach calculates contribution weights for each node’s relevance to degradation and applies weighted pooling accordingly. The PROTAC graph embedding 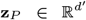 is computed as follows:

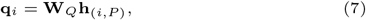

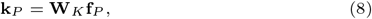

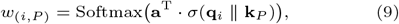

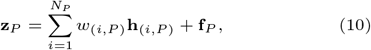

where 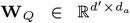 and 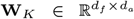 are projection matrices, with *d*^*′*^, *d*_*f*_, and *d*_*a*_ representing the feature dimensions of node embeddings, selected molecular attributes, and attention, respectively. The vector 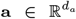 serves as the attention vector, and σ(·) is the activation function, where we utilise tanh(·). Using the same procedure, the graph embeddings **z**_*T*_ and **z**_*E*_ for POI and E3 ligase graphs are computed. The final representation **z** for the POI-PROTAC-E3 ligase ternary complex is obtained by concatenating the three graph embeddings.

#### 2.2.5. Label enrichment

As discussed in Section 2.1, only 1,502 samples out of the total 8,603 PROTAC entries in the collected dataset are labelled. Thus, employing a semi-supervised learning approach that integrates a large volume of unlabeled data into the training process is advantageous for addressing the lack of annotated samples and enhancing prediction accuracy [7]. Inspired by recent advancements in pseudo-labelling based on training dynamics [31], we propose a memory-enhanced pseudo-labelling strategy to generate and select pseudo-labels using a combined selection score. Specifically, we adopt the disagreement score design from MoDis [31] and propose a novel memory-based score.

Following the process proposed in [31], we compute the disagreement score by first pre-training the DegradeMaster model on labelled samples to predict the labels of unlabeled data. As illustrated in Figure 1(D), for each unlabeled entry *k*, the predicted labels across selected training epochs form the pseudo-label set ℳ. The logits **z**_*k*_ are used to compute the prediction distribution *P*_*k*_(*c*), reflecting the relative frequency of class *c*. The disagreement score *s*_*dis*_(*k*), calculated as the cross-entropy of *P*_*k*_(*c*), measures the consistency of predictions for *k*. A higher *s*_*dis*_(*k*) indicates a stronger candidate for pseudo-labeling.

To refine pseudo-label selection further, we design a memory-based score using latent feature similarity. We record the latent embedding 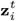 for each labeled entry *i* ∈ *U*_*l*_ at epoch *t* ∈ 𝒯, and calculate the memory prototype 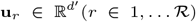 for the labeled data. Here, ℛ is a hyper-parameter determining the number of prototypes, and **u**_*r*_ is computed as:

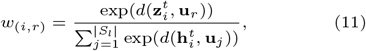

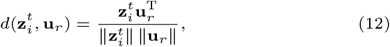

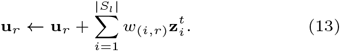

The latent embedding of the unlabeled entry *k* ∈ *S*_*u*_ at epoch *t* ∈ 𝒯 is denoted as 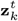. The memory-based score *s*_*mem*_(·) is calculated as the cosine similarity between 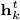 and the memory prototypes **u**_*r*_:

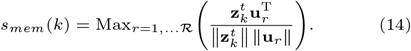

The *s*_*mem*_(·) score measures the similarity between the latent embedding of unlabeled data and labelled data. Higher *s*_*mem*_ values indicate more promising pseudo-labeling candidates. By combining *s*_*dis*_(·) and *s*_*mem*_(·), we acquire the combined selection score *s*:

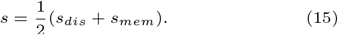

After 𝒯 training epochs, we rank the unlabeled data based on *s* score and select the top *K* samples with the highest scores. The corresponding predicted labels are chosen from ℳ as pseudo-labels, which are then combined with real labels to re-train DegradeMaster. Note that we only use the training set for label enrichment and directly apply the re-trained model for prediction on the test set.

## 3. Results and Discussion

We use information from the PROTAC-DB 3.0 [14] as described in 2.1, and construct a supervised PROTAC dataset named PROTAC-1K and a semi-supervised PROTAC dataset named PROTAC-8K. We randomly split both PROTAC-1K and PROTAC-8k datasets into training/testing sets with an 80/20 ratio. Note that PROTAC-1K and PROTAC-8K share identical labelled data in both their training and testing sets. We trained SVM and RF in a supervised setting using PROTAC-1K and trained three GNN-based models in both supervised and semi-supervised settings. Model configuration is shown in Table S3 in supplementary. The performance of DegradeMaster was evaluated against four baselines: support vector machine (SVM) [15], random forest (RF) [16], DeepPROTACs [25], and PROTAC-STAN [24] under the same random split setting, using the metrics outlined in Section 2 in supplementary. For a fair comparison, we apply the same random seed for all models.

### 3.1. Model performance

The comparison results are presented in Table 2, with corresponding ROC curves shown in Figure 3. Both SVM and RF methods outperform DeepPROTACs across all three metrics. Specifically, SVM achieves a 15.76% improvement in accuracy, while RF shows a 13.64% higher AUROC score compared to DeepPROTAC. These results underscore the efficacy of machine learning approaches, even in their simplicity, particularly when applied to limited training data. Moreover, the fingerprint-based features prove effective, with Morgan fingerprints demonstrating a slight advantage over MACCS fingerprints. Consequently, we selected Morgan fingerprints and six additional physico-chemical properties (elaborated in Section S1) to construct the raw features for three deep graph learning models. The results with and without the raw features are depicted in Figure 2.

**Table 2.** The evaluation results of DegradeMaster and the baseline methods on the test set. The SVM and RF methods are implemented with two fingerprints, i.e., MACCS and Morgan. In all metrics, DegradeMaster (bold) outperformed alternative models, while the best-performing baselines are underlined. Abbreviation: Accuracy (Acc), Fingerprints (FGP), Supervised (Super), Semi-supervised (Semi).

| Method | FGP | Setting | Acc (%) | AUROC | F1 score |
| --- | --- | --- | --- | --- | --- |
| SVM | *MACCS* | *Super* | 72.34 | 0.7815 | 0.7232 |
|  | *Morgan* | *Super* | 74.47 | 0.7901 | <u>0.7446</u> |
| RF | *MACCS* | *Super* | 67.65 | 0.7981 | 0.6498 |
|  | *Morgan* | *Super* | 63.40 | 0.8077 | 0.5818 |
| DeepPROTACs | - | *Super* | 58.21 | 0.7107 | 0.5714 |
|  | - | *Semi* | 59.02 | 0.7171 | 0.5731 |
| PROTAC-STAN | - | *Super* | <u>79.34</u> | <u>0.7986</u> | 0.7310 |
|  | - | *Semi* | 78.25 | 0.7896 | 0.7403 |
| DegradeMaster | - | *Super* | 81.41 | 0.8541 | 0.8139 |
|  | - | *Semi* | **83.66** | **0.8825** | **0.8224** |

**Fig. 2.**
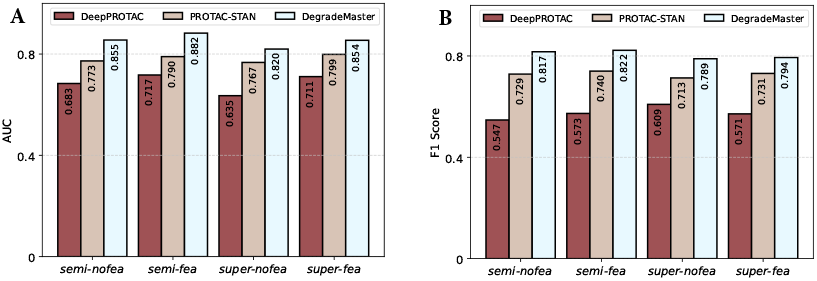
Performance comparison between DegradeMaster and two deep learning baselines (DeepPROTAC and PROTAC-STAN) on the testing dataset: (**A**) AUC comparison of models under four different settings. (**B**) F1 comparison of models under four settings. Abbreviation: semi-supervised with no features (semi-nofea), semi-supervised with features (semi-fea), supervised with no features (super-nofea), supervised with features (super-fea)

**Fig. 3.**
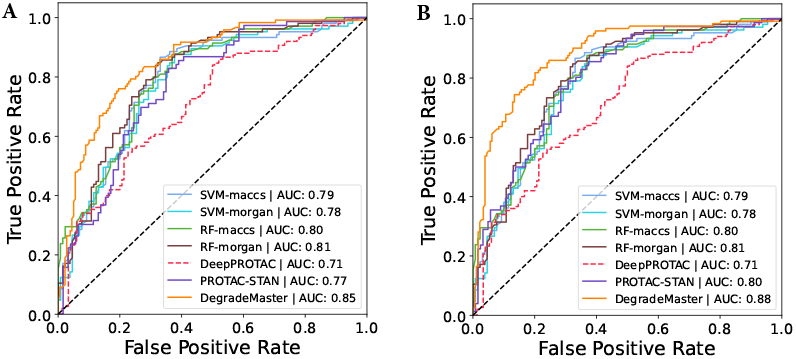
ROC curves of DegradeMaster and compared baselines. (**A**) ROC curves on PROTAC-1K. (**B**) ROC curves on PROTAC-8K.

Both variants of DegradeMaster consistently surpass the existing methods, demonstrating superior performance across accuracy, AUROC, and F1 score. This success is attributed to the E(3)-equivariant design of the graph encoder, which effectively integrates the spatial and structural properties of molecules in 3D space into the representations, thereby enhancing downstream prediction tasks. Specifically, DegradeMaster surpasses the previous state-of-the-art model, PROTAC-STAN, by 2.6% in accuracy, 6.9% in AUROC, and 11.34% in F1 score under the supervised setting. Additionally, it achieves a 6.9% improvement in accuracy, 11.76% in AUROC, and 8.21% in F1 score under the semi-supervised setting. These results underscore DegradeMaster’s advanced predictive capabilities and its effectiveness in PROTAC degradation prediction. Furthermore, comparing the performance of the supervised and semi-supervised variants of the three deep learning models reveals contrasting trends. Existing models like DeepPROTACs exhibit only marginal performance gains when additional unlabeled PROTAC entries are incorporated, while PROTAC-STAN experiences slight performance degradation. These results demonstrate that incorporating unlabeled data without carefully designed mechanisms for label enhancement does not necessarily improve model generalization or performance. In contrast, DegradeMaster demonstrates a notable improvement of 3.3% in AUROC under the semi-supervised setting. This underscores the effectiveness of our label enhancement strategy, which identifies high-quality unlabeled data and generates pseudo-labels for inclusion in the training set. By addressing the challenge of label scarcity, this approach enhances the scalability and performance of our model.

### 3.2. Ablation Study

In this section, we conduct an ablation study to analyze the contributions of the core components of DegradeMaster, as described in Section 2.2. Table 3 summarizes the results. Five variants of DegradeMaster are evaluated, including four degraded versions and the full model. Each variant isolates specific modules to assess their individual and collective impact. The variants are described as follows:

**Table 3.** Ablation study. DM is the abbreviation of DegradeMaster.

| Component | Feature Selection | Attention pooling | Equivariant encoder | Label enrichment | Acc (%) | AUROC | F1 score |
| --- | --- | --- | --- | --- | --- | --- | --- |
| DM<sub>raw</sub> | ✗ | ✗ | ✗ | ✗ | 70.33 | 0.8104 | 0.6987 |
| DM<sub>w/o feat</sub> | ✗ | ✓ | ✓ | ✓ | 78.33 | 0.8583 | 0.7720 |
| DM<sub>w/o att</sub> | ✓ | ✗ | ✓ | ✓ | 76.33 | 0.8229 | 0.7630 |
| DM<sub>w/o e(3)</sub> | ✓ | ✓ | ✗ | ✓ | 72.66 | 0.8044 | 0.7201 |
| DM<sub>w/o pseu</sub> | ✓ | ✓ | ✓ | ✗ | 80.01 | 0.8575 | 0.7891 |
| DM<sub>full</sub> | ✓ | ✓ | ✓ | ✓ | 83.66 | 0.8825 | 0.8224 |

- DegradeMaster_*w/o feat*_: This variant removes the feature selection module, using all chemicophysical properties from PROTAC-DB 3.0 as raw input features.
- DegradeMaster_*w/o att*_: This variant excludes the attention mechanism by replacing attention pooling with mean pooling.
- DegradeMaster_*w/o e*(3)_: This version substitutes the E(3)-equivariant encoder with a standard GCN [19] encoder, removing the 3D spatial structural representation.
- DegradeMaster_*w/o pseu*_: This variant disables the label enrichment strategy by setting *K* to 0, thereby omitting pseudo-labeling.
- DegradeMaster_*raw*_ : This baseline excludes all key components, including feature selection, attention pooling, the E(3)-equivariant encoder, and label enrichment.
- DegradeMaster_*full*_: Full model of DegradeMaster with all the components available.

From Table 3, it is evident that DegradeMaster_*raw*_ demonstrates a significant performance decline compared to DegradeMaster_*full*_, with 18.95% decrease in accuracy and 17.7% in F1 score. This underscores the importance of the overall design and the integration of the proposed key components. When comparing DegradeMaster_*w/o e*(3)_ with DegradeMaster_*full*_, the largest performance reduction of removing a single component is observed, highlighting the critical role of the E(3)-equivariant encoder in degradation prediction tasks. This finding emphasizes the necessity of incorporating 3D spatial and structural information of proteins and molecules into the representation. The removal of the attention pooling module (DegradeMaster_*w/o att*_) leads to a 9.6% decrease in accuracy and a 7.24% reduction in AUROC compared to DegradeMaster_*full*_. This result demonstrates the effectiveness of the proposed attention mechanism in selectively aggregating node embeddings in proteins and molecules. The elimination of the pseudo-labeling component results in the reduction of accuracy by 4.56% and AUROC by 2.91%. This demonstrates the efficacy of the proposed pseudo-labeling strategy in leveraging the rich information from unlabeled data.

### 3.3. Attention weight visualization

As described in Section 2.2.4, we implement a mutual attention mechanism during pooling to compute protein-level and molecule-level embeddings. To facilitate interpretability, we visualize the attention weights by mapping them back onto the atom-level representation of the PROTAC molecule and the amino acid residue-level representation of the E3 ligase and target protein. This enables a bioinformatic analysis of the significance and distribution of the attention values. Figure 4 illustrates the attention weights among atoms within a PROTAC molecule [32] that facilitates the degradation of the BRD2 protein via the CRBN ligase. The 3D molecular graphs are constructed from the atomic coordinates provided in the mol2 file, with edge colour intensity reflecting the attention weights among atoms. Based on Figure 4, we derive the following observations:

**Fig. 4.**
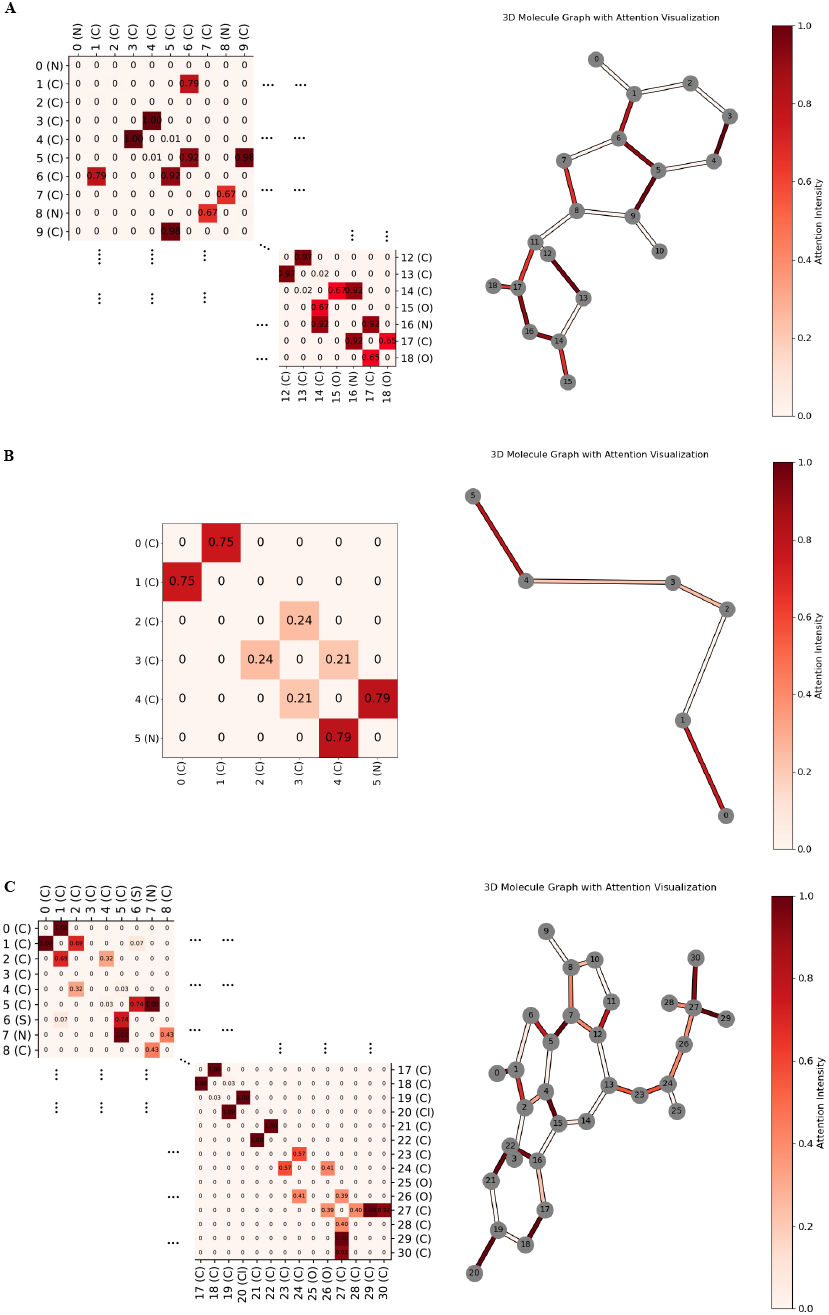
Visualization of attention weights for the PROTAC molecule (PROTAC-DB ID: 194) facilitating degradation of the BRD2 target protein via the CRBN E3 ligase. The left panel displays an attention heatmap among atoms within the molecule, while the right panel presents the 3D molecular graph, where edge color intensity represents attention values. (**A**) Attention values targeting the E3 ligand. (**B**) Attention values are associated with the linker. (**C**) Attention values targeting the warhead.

Linking areas show higher attention weights: Atoms located at the interface regions connecting the warhead- and E3-binding ligand exhibit higher attention weights. For instance, in Figure 4(A), the edge between atom 1 (C) and atom 6 (C) has an attention weight of 0.794, while the edge between atom 6 (C) and atom 5 (C) has an attention weight of 0.917. Similarly, in Figure 4(C), the edges between atom 27 (C) and atoms 28 (C), 29 (C), and 30 (C) have significantly higher weights compared to the central region, such as between atom 13 and atom 14. This highlights the model’s ability to recognise linking regions of target/E3 ligands and emphasizes the importance of structural information in these areas for degradation prediction.

Ends of the linker have higher attention: As observed in Figure 4(B), atoms at both ends of the linker display higher attention weights than those in the middle. This aligns with the actual interaction modes between the linker and the target/E3 ligands.

Binding regions of the POI target and E3 Ligands show higher attention: Atoms at the distal ends of the target and E3 ligands also exhibit elevated attention weights. For example, in Figure 4(C), atoms 20 (Cl) and 19 (C), as well as atoms 18 (C) and 17 (C), show higher attention weights. Similarly, in Figure 4(A), atoms 17 (C) and 16 (N), as well as atoms 16 (N) and 14 (C), demonstrate higher weights. As depicted in Figure S1 in supplementary, these atoms are located near the binding pockets of the POI and E3 ligase (e.g., GLU-187 in the E3 ligase and GLU-735 in the POI). These patterns correspond to the actual binding modes [33, 34], with the warhead binding to the target protein and the E3 ligand interacting with the E3 ligase.

Figure S1 illustrates the binding interactions between the PROTAC molecule and protein pockets. The cyan region represents the CRBN ligase, while the purple region denotes the POI. PROTAC is shown in green. Residues located within 5 Å of the ligase ligand and warhead are selected as the binding pocket of E3 ligase and POI, respectively. The attention weight for each residue is calculated as the mean of the attention weights of the atoms within that residue. Residues with the highest attention weights are labelled in Figure S1 (A), with their corresponding attention values displayed in Figures S1 (B) and S1 (C). Notably, residues PRO-186 and GLU-187 in the ligase pocket, as well as residues GLU-735 and LYS-732 near the binding regions of the ligase ligand and warhead, are identified as high-weight residues, highlighting their importance in the binding interaction.

## 4. Case Study

To further demonstrate the predictive capabilities of DegradeMaster, we conduct two case studies. Case Study 1 focuses on predicting candidates during the development of the PROTAC VZ185 [9], while Case Study 2 evaluates the model’s ability to predict the degradability of PROTAC ACBI3 across different KRAS mutants [35].

### 4.1. Case study 1

In Case Study 1, we constructed an experimental dataset comprising 16 PROTAC candidates developed during the design of PROTAC VZ185 [9], which recruits VHL ligase to degrade BRD9, a chromatin binding factor that plays a crucial role in regulating gene expression. The dataset includes various linear alkyl chains and polyethylene glycol (PEG) chains of different lengths as linkers, paired with two variants of the VHL E3 ligase binding ligand (VHL3 and VHL4), resulting in 16 PROTAC candidates. As shown in Figure 5, the rightmost column indicates the actual performance of these PROTAC candidates. Of the 16, five exhibit high BRD9 degradation, while the remaining 11 have relatively low efficacy. Figure 6 (A) illustrates that DegradeMaster achieves 83.33% accuracy and an AUROC of 0.9692 in predicting the BRD9 degradation by these PROTAC candidates, significantly outperforming two baseline models. Notably, as shown in Figure 5, DegradeMaster demonstrates accurate predictions for both positive and negative samples, with only two false-positive predictions. These results highlight the potential of DegradeMaster to accelerate PROTAC development by nominating and screening promising candidates, thereby streamlining iterative the drug discovery process.

**Fig. 5.**
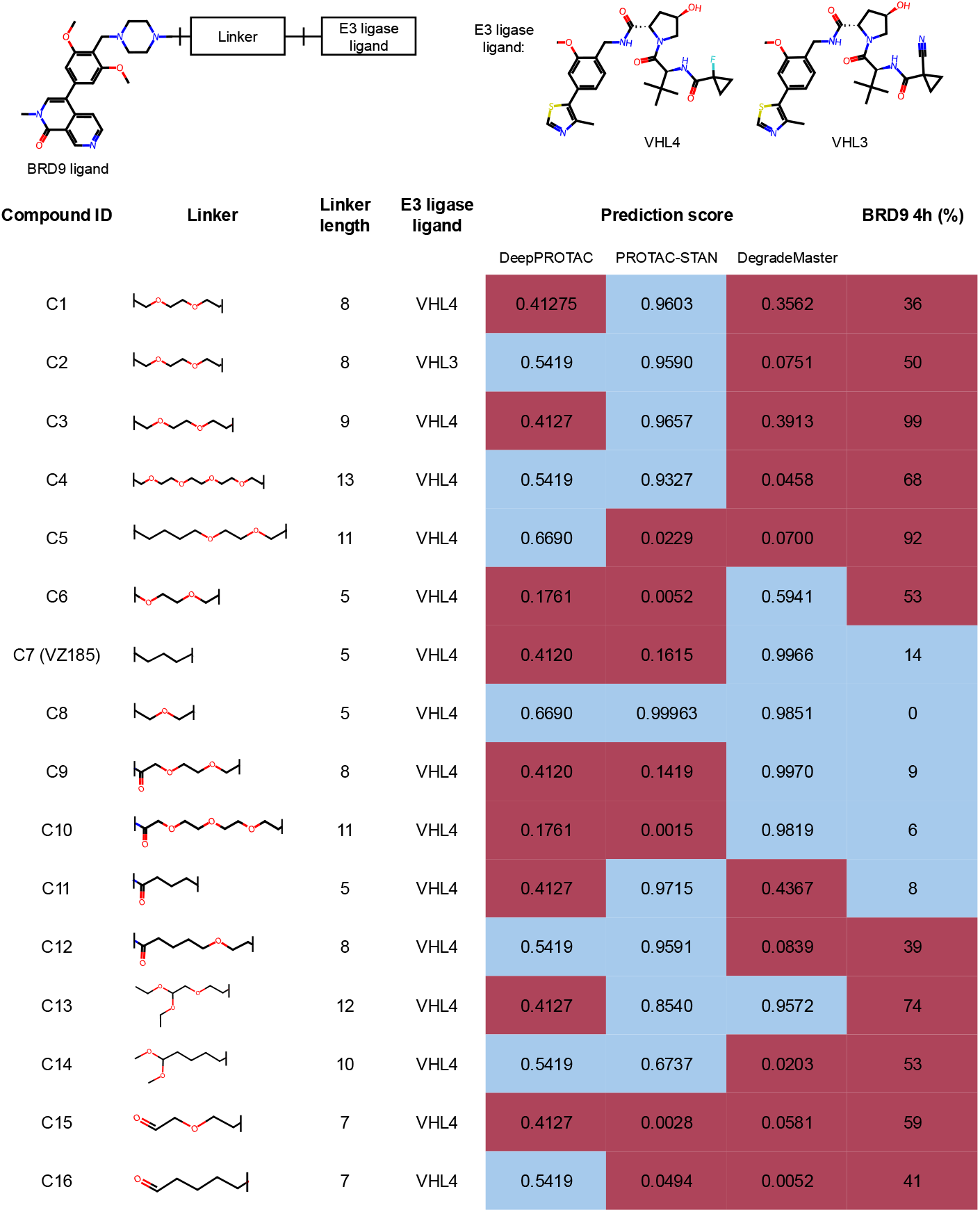
Case study 1: degradability prediction of the VZ185 [9] PROTAC candidates. Compounds 13–16 only consist of a linker and an E3 binder, with no target binder included. The rightmost column represents the degradation activity, expressed as the percentage of total protein remaining after treatment with 1*μM* compound. Prediction scores range from 0 to 1, with higher scores indicating a greater likelihood of protein degradation. Threshold of high degradation (blue) and low degradation (red): 0.5 for prediction scores and 15% for protein remaining.

**Fig. 6.**
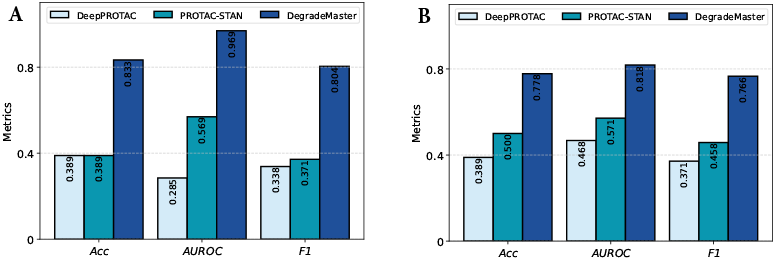
Performance comparison of DegradeMaster associated with two DL models on case study 1 (**A**) and case study 2 (**B**).

### 4.2. Case study 2

Mutations in the Kirsten rat sarcoma viral oncogene homolog (KRAS) GTPase protein are among the most prevalent in cancer. Developing small molecule inhibitors or PROTACs capable of targeting oncogenic KRAS alleles remains challenging, with most progress limited to KRAS with a glycine to cysteine mutation at position 12 (G12C), for which specific small molecules with covalent warheads have been developed. Recently, a study [35] successfully developed a ‘universal’ pan KRAS degrader, ACBI3, a heterobifunctional small molecule that utilises a noncovalent KRAS ligand to bind and potently. degrade 13 out of the 17 most prevalent oncogenic KRAS alleles [35].

In Case Study 2, we evaluated the degradability of KRAS by ACBI3 [35] on 17 mutants, presented in Figure 6 and Figure S2. DegradeMaster demonstrates superior performance across all three metrics compared to two baseline models. Notably, since the only variable among the 17 samples is the residue point mutation present in the POI sequence, these results highlight DegradeMaster’s capability to effectively capture both structural and sequence information of the POI. This further validates the potential of DegradeMaster to drive the development of novel PROTAC candidates against even highly mutagenised POI targets.

## 5. Model Efficiency

This section compares the computational efficiency of DegradeMaster with two existing models. For a fair comparison, we set the hidden dimension as 128 and batch size as 50 for all three models. Figure 7 shows that DegradeMaster has similar training time to DeepPROTAC for 1,000 samples but slightly higher inference time. As dataset size grows, DeepPROTAC is 1.2 × faster and also consumes less memory usage (Table 4), due to its simpler 2D graph modeling and GCN encoding design. However, this simplicity comes at the cost of predictive performance, with DeepPROTAC showing a 23% lower AUROC compared to DegradeMaster. Compared to PROTAC-STAN, DegradeMaster is 1.2 × faster in training, uses 27% less memory for training, and 86% less for inference. Additionally, DegradeMaster scales more efficiently with dataset size due to its lightweight mutual attention mechanism, avoiding the computational overhead of PROTAC-STAN’s ternary attention networks.

**Table 4.** Comparison of training and inference memory usage across three models: DeepPROTAC, PROTAC-STAN and DegradeMaster.

| Mode | PROTAC-STAN | DeepPROTAC | Degrademaster |
| --- | --- | --- | --- |
| Training | 31,277MB | 4,081MB | 22,757MB |
| Inference | 11,513MB | 1,509MB | 1,613MB |

**Fig. 7.**
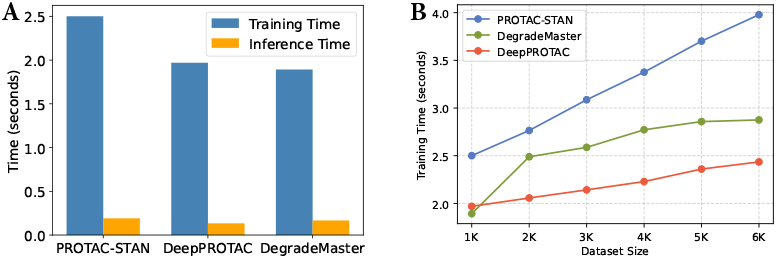
**(A)** Training and inference time per epoch for DeepPROTAC, PROTAC-STAN and DegradeMaster with 1K of training samples. **(B)** The curve of training time when varying the dataset size from 1K to 6K.

## 6. Conclusion

This study introduces DegradeMaster, a semi-supervised E(3)-equivariant graph neural network designed to predict PROTAC-targeted degradation with enhanced performance.

By integrating 3D geometric constraints through an E(3)-equivariant encoder, employing a memory-based pseudo-labeling for label enrichment, and utilising a mutual attention pooling module for comprehensive graph embedding, DegradeMaster addresses the limitations of spatial information loss and label scarcity in existing approaches. Experiments demonstrate its superior performance over state-of-the-art methods and high accuracy in case studies involving challenging degradability predictions. These results highlight the critical role of 3D structural information and semi-supervised learning in advancing computational methods for PROTAC degradation prediction.

## Notes

https://zenodo.org/records/14728925

https://github.com/Jackson117/DegradeMaster

### Competing Interest Statement

The authors have declared no competing interest.

