## Supplementary for "Accurate PROTAC targeted degradation prediction with DegradeMaster"

### 1. FEATURE SELECTION

In addition to spatial and structural information, the intrinsic physicochemical properties of molecules can provide valuable insights into degradation. This section introduces our feature selection pipeline to identify the most informative chemical properties. As shown in Table S1, we extract 11 chemical properties of PROTAC molecules from PROTAC-DB 3.0. Directly inputting all raw attributes into the deep learning model risks introducing noise from irrelevant features, potentially reducing predictive performance. To address this, we implement a feature selection module with two steps, using only training set data. The first step employs a filter selection approach based on statistical metrics, including variance, Pearson correlation, chi-square score, p-value, and mutual information score, defined as follows:

- **Variance**: A metric measuring the spread of feature values from the mean. Features with very low variance are often irrelevant for distinguishing between target classes. Here we set the filtering threshold as 0.01.

- **Pearson Correlation Coefficient**: A metric measuring the strength and direction of a linear relationship between two variables, ranging from -1 (perfect negative relationship) to 1 (perfect positive relationship), with 0 indicating no linear relationship. The threshold is set as $\pm 0.9$.

- **Chi-Square Score**: A statistical test used to evaluate the independence between features and the label. Higher scores indicate stronger associations. We set the filtering threshold as 10.

- **P-Value**: A metric that assesses the significance of a feature in hypothesis testing. Lower p-values suggest that the feature is significantly related to the label. The threshold is set as 0.05.

- **Mutual Information**: A metric that measures the amount of information shared between the feature and the label. Higher mutual information indicates a stronger relationship.

As shown in Table S1, the variance of Gasteiger charges is significantly lower than that of other properties, leading to its exclusion. The correlation matrix for all 11 molecular properties is presented in Table S2, revealing that exact mass and heavy atom count are highly correlated with molecular weight. Consequently, these two properties are filtered as redundant. Based on the chi-square score and p-value columns, we also exclude hydrogen bond acceptor count and chiral centres. However, most properties exhibit low mutual information scores, suggesting that mutual information may not be a reliable metric for feature selection in this context.

Following the filter method, we apply an embedded method for further refinement. Specifically, we use Gradient Boosting Decision Trees (GBDT) [1] as the base model, training it with the filtered features and labels to compute feature importance as the normalized total reduction in the splitting criterion. Table S1 reports the results, where hydrogen bond donor count and rotatable bond count show the lowest importance. Nevertheless, given their high chi-square scores, these descriptors are retained, underscoring the importance of integrating multiple statistical metrics in feature selection.

In conclusion, the two-step feature selection process resulted in the preservation of six chemical properties: molecular weight, XLogP3, ring count, hydrogen bond donor count, rotatable bond count, and topological polar surface area as the final selected features for PROTAC molecules. For the POI and E3 ligase proteins, we adopt the auto-cross covariance (ACC) method, as described

| **Chemical properties** | **Variance** | **Chi-square score** | **P-value** | **Mutual-info score** | **GBDT importance** | **Selection** |
|---|---|---|---|---|---|---|
| Molecule weight | 28345.75 | 10.04 | 0.0015 | 0.0720 | 0.2260 | ✓ |
| Exact mass | 28287.18 | 10.08 | 0.0014 | 0.0950 | – | |
| XLogP3 | 5.96 | 13.77 | 0.0002 | 0.0778 | 0.2029 | ✓ |
| Heavy Atom Count | 130.56 | 0.1821 | 0.6695 | 0.0186 | – | |
| Ring Count | 1.83 | 20.63 | ≈0 | 0.0537 | 0.1928 | ✓ |
| Hydrogen Bond Acceptor Count | 8.85 | 0.0650 | 0.7987 | 0.0226 | – | |
| Hydrogen Bond Donor Count | 2.87 | 26.95 | ≈0 | 0.0055 | 0.0556 | ✓ |
| Rotatable Bond Count | 40.71 | 70.78 | ≈0 | 0.0071 | 0.0551 | ✓ |
| Topological Polar Surface Area | 2004.48 | 340.9 | ≈0 | 0.1764 | 0.2673 | ✓ |
| Gasteiger charges | 0.01 | 0.0034 | 0.9533 | 0.1844 | – | |
| Chiral centers | 3.47 | 1.241 | 0.2651 | 0.0133 | – | |

**Table S1.** Chemical properties of PROTAC molecules collected from PROTAC-DB 3.0 and their corresponding statistical metrics.

| | Molecular Weight | Exact Mass | XLogP3 | Heavy Atom Count | Ring Count | Hydrogen Bond Acceptor Count | Hydrogen Bond Donor Count | Rotatable Bond Count | Topological Polar Surface Area | Charges | Chiral Centers |
|---|---|---|---|---|---|---|---|---|---|---|---|
| Molecular Weight | nan | **0.9999** | 0.4694 | **0.9871** | 0.3956 | 0.6233 | 0.4244 | 0.7538 | 0.6047 | 0.2627 | 0.5167 |
| Exact Mass | nan | nan | 0.4691 | **0.9873** | 0.3958 | 0.6236 | 0.4246 | 0.7539 | 0.6051 | 0.2627 | 0.5167 |
| XLogP3 | nan | nan | nan | 0.4620 | 0.2595 | -0.1457 | -0.0051 | 0.2262 | -0.2374 | 0.2851 | 0.3898 |
| Heavy Atom Count | nan | nan | nan | nan | 0.4677 | 0.6283 | 0.4029 | 0.7376 | 0.6061 | 0.2557 | 0.5086 |
| Ring Count | nan | nan | nan | nan | nan | 0.2880 | -0.2365 | -0.0587 | 0.0424 | 0.0365 | 0.0882 |
| Hydrogen Bond Acceptor Count | nan | nan | nan | nan | nan | nan | 0.2694 | 0.6372 | 0.7310 | 0.1547 | 0.1128 |
| Hydrogen Bond Donor Count | nan | nan | nan | nan | nan | nan | nan | 0.4980 | 0.7296 | -0.0199 | 0.3532 |
| Rotatable Bond Count | nan | nan | nan | nan | nan | nan | nan | nan | 0.6444 | 0.1370 | 0.3203 |
| Topological Polar Surface Area | nan | nan | nan | nan | nan | nan | nan | nan | nan | -0.0195 | 0.2277 |
| Charges | nan | nan | nan | nan | nan | nan | nan | nan | nan | nan | 0.2477 |
| Chiral Centers | nan | nan | nan | nan | nan | nan | nan | nan | nan | nan | nan |

**Table S2.** Correlation matrix for molecular descriptors, computed using pearson correlation method. The correlations above the threshold 0.9 are in bold.

in [2], to represent protein sequence features. ACC quantifies the relationship between amino acid properties within a protein sequence across varying lags.

### 2. PERFORMANCE EVALUATION METRICS

To assess the performance of DegradeMaster and its competitors, we utilize a combination of ROC-AUC, precision, recall, and Macro-F1 as evaluation metrics. ROC-AUC, a standard metric in anomaly detection, involves plotting the true positive rate against the false positive rate, with AUC (Area Under the Curve) representing the area under this ROC curve, ranging between 0 and 1. Higher AUC values indicate better performance. Additionally, we use precision, recall, and macro-F1 as our evaluation metrics. For each class, the precision, recall, and $F_1$ score can be calculated as follows:

$$Precision = \frac{TP}{TP + FP},$$

$$Recall = \frac{TP}{TP + FN},$$

$$F_1 = 2 \times \frac{Precision \times Recall}{Precision + Recall},$$

where $TP$ denotes the number of positive samples that are correctly predicted; $FP$ is the number of negative samples that are predicted as the positive ones; and $FN$ is the number of positive samples that are predicted as the negative ones. Macro-F1, as the harmonic mean of precision and recall, is computed individually for each class and then averaged, to provide a balanced measure of the model's performance. Therefore, by averaging the $F_1$ scores of both classes, macro-F1 can be obtained as follows:

$$Macro\text{-}F1 = \frac{F_1(L) + F_1(H)}{2},$$

where $L$ and $H$ denote the low-activity and the high-activity class, respectively.

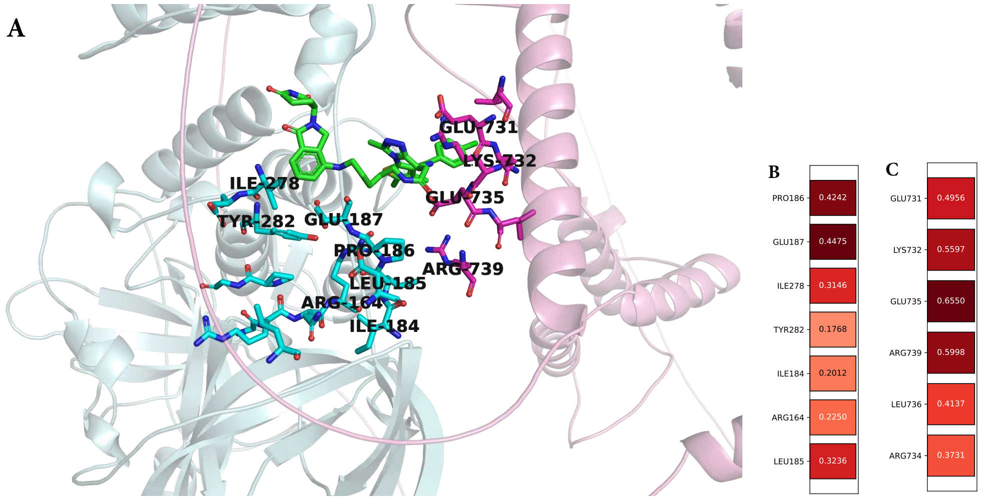


**Fig. S1.** (**A**) Visualization of the ternary complex comprising the PROTAC molecule (PROTAC-DB ID: 194), POI, and E3 ligase. The E3 ligase is shown in cyan, and the POI is depicted in light pink. The 5 Å protein pockets of both the E3 ligase and POI are further visualized, with residue names labeled. The PROTAC molecule is represented in green. (**B**) Attention weight distribution for residues within the E3 ligase pocket. (**C**) Attention weight distribution for residues within the POI pocket. The attention weight of each residue is computed as the average of the attention weights assigned to all atoms within the residue.

| Mutant ID | Mutant Sequence | Mutation | PROTAC | Prediction | | | Degradability |
|---|---|---|---|---|---|---|---|
| | | | | DeepPROTAC | PROTAC-STAN | Ours | |
| 1 | . . . V V V G A G G V G K S . . . | None | ACBI3 | 0.3172 | 0.9876 | 0.9379 | $DC_{50} \leq 100nM$ |
| 2 | . . . V V V G A **C** G V G K S . . . | G12C | ACBI3 | 0.5419 | 0.9743 | 0.1936 | $DC_{50} \leq 100nM$ |
| 3 | . . . V V V G A **A** G V G K S . . . | G12A | ACBI3 | 0.3168 | 0.9875 | 0.9382 | $DC_{50} \leq 100nM$ |
| 4 | . . . V V V G A **D** G V G K S . . . | G12D | ACBI3 | 0.5419 | 0.9752 | 0.1945 | $DC_{50} \leq 100nM$ |
| 5 | . . . V V V G A **V** G V G K S . . . | G12V | ACBI3 | 0.6690 | 0.9998 | 0.9801 | $DC_{50} \leq 100nM$ |
| 6 | . . . V V G A G **C** V G K S A . . . | G13C | ACBI3 | 0.6690 | 0.9998 | 0.9801 | $DC_{50} \leq 100nM$ |
| 7 | . . . V V G A G **D** V G K S A . . . | G13D | ACBI3 | 0.6361 | 0.0047 | 0.9982 | $DC_{50} \leq 100nM$ |
| 8 | . . . V V G A G **V** V G K S A . . . | G13V | ACBI3 | 0.1761 | 0.0067 | 0.8824 | $DC_{50} \leq 100nM$ |
| 9 | . . . L D T A G **E** E E Y S A . . . | Q61E | ACBI3 | 0.2077 | 0.1675 | 0.9946 | $DC_{50} \leq 100nM$ |
| 10 | . . . F I E T S **V** K T R Q R . . . | A146V | ACBI3 | 0.1761 | 0.0069 | 0.8828 | $DC_{50} \leq 100nM$ |
| 11 | . . . L D T A G **P** E E Y S A . . . | Q61P | ACBI3 | 0.3180 | 0.9879 | 0.9374 | $DC_{50} \leq 100nM$ |
| 12 | . . . F I E T S **P** K T R Q R . . . | A146P | ACBI3 | 0.3172 | 0.9876 | 0.9384 | $DC_{50} \leq 100nM$ |
| 13 | . . . F I E T S **T** K T R Q R . . . | A146T | ACBI3 | 0.5419 | 0.9728 | 0.0352 | $DC_{50} \leq 100nM$ |
| 14 | . . . L D T A G **H** E E Y S A . . . | Q61H | ACBI3 | 0.5419 | 0.1222 | 0.1922 | $DC_{50} > 100nM$ |
| 15 | . . . L D T A G **K** E E Y S A . . . | Q61K | ACBI3 | 0.3163 | 0.9766 | 0.9379 | $DC_{50} > 100nM$ |
| 16 | . . . L D T A G **L** E E Y S A . . . | Q61L | ACBI3 | 0.5196 | 0.9875 | 0.0323 | $DC_{50} > 100nM$ |
| 17 | . . . L D T A G **R** E E Y S A . . . | Q61R | ACBI3 | 0.5419 | 0.1190 | 0.1958 | $DC_{50} > 100nM$ |
| 18 | . . . V V V G A **R** G V G K S . . . | G12R | ACBI3 | 0.5420 | 0.9743 | 0.1936 | $DC_{50} > 100nM$ |

**Fig. S2.** Case study 2: degradation prediction of the ACBI3 [3] on KRAS mutants. Protein 1 has no mutations whereas the others have a mutated residue at the corresponding position. The rightmost column represents the degradation activity, expressed as the $DC_{50}$ values. Prediction scores range from 0 to 1, with higher scores indicating a greater likelihood of protein degradation. Threshold of high degradation (blue) and low degradation (red): 0.5 for prediction scores and 100 nM for $DC_{50}$.

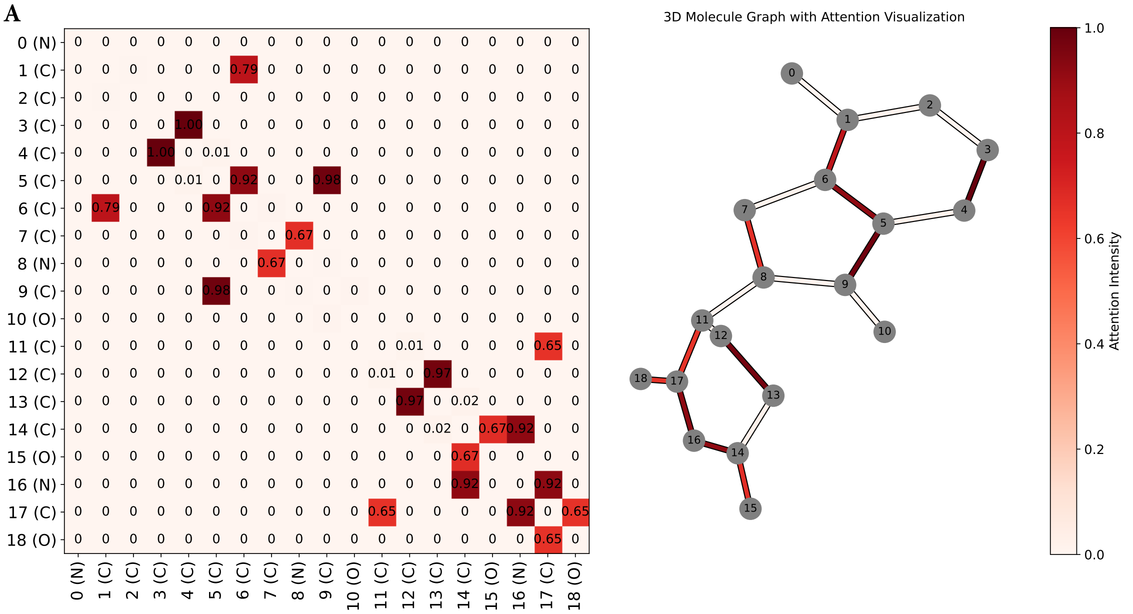


**Fig. S3.** Visualization of attention weights for the E3 ligand of PROTAC molecule (PROTAC-DB ID: 194), including the complete attention matrix.

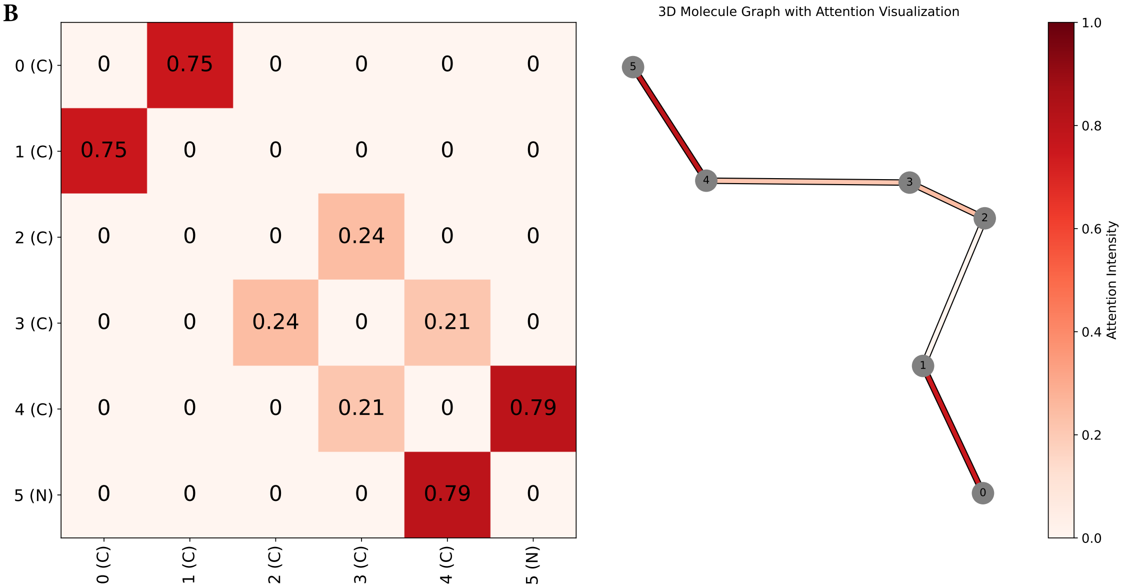


**Fig. S4.** Visualization of attention weights for the linker of PROTAC molecule (PROTAC-DB ID: 194), including the complete attention matrix.

C

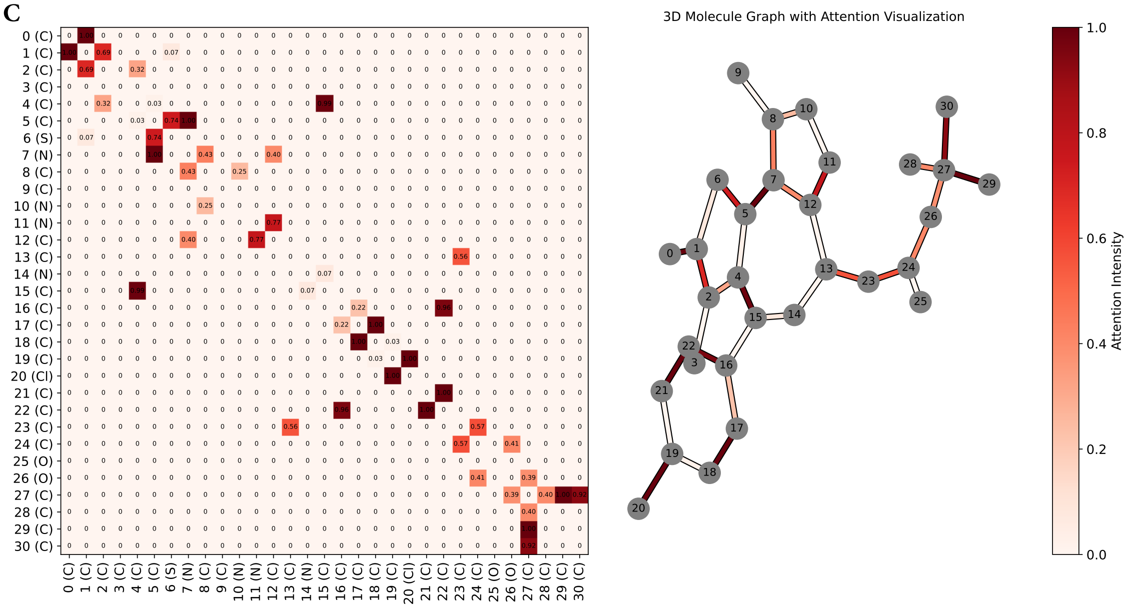


**Fig. S5.** Visualization of attention weights for the warhead of PROTAC molecule (PROTAC-DB ID: 194), including the complete attention matrix.

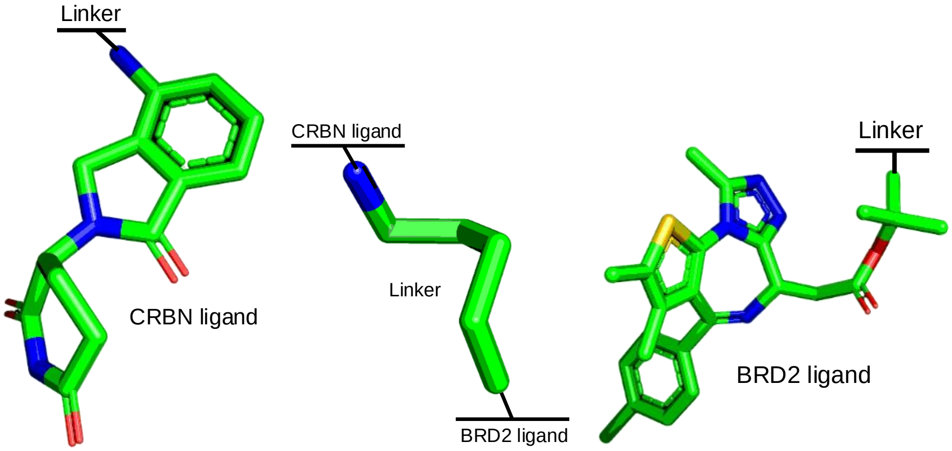


**Fig. S6.** The chemical structures of the E3 ligand, linker, and warhead of PROTAC-194.
